# *C. elegans* Mediator 15 permits low temperature-induced longevity via regulation of lipid and protein homeostasis

**DOI:** 10.1101/366153

**Authors:** Dongyeop Lee, Seon Woo A. An, Yoonji Jung, Yasuyo Yamaoka, Youngjae Ryu, Grace Ying Shyen Goh, Arshia Beigi, Dengke Ma, Chang Man Ha, Stefan Taubert, Youngsook Lee, Seung-Jae V. Lee

## Abstract

Low temperatures slow aging and extend lifespan in many organisms, including *Caenorhabditis elegans.* However, the metabolic and homeostatic aspects of low temperature-induced longevity remain poorly understood. Here, we show that changes in lipid composition regulated by MDT-15/Mediator 15, transcriptional co-regulator, is essential for low temperature-induced longevity and proteostasis in *C. elegans.* We find that inhibition of *mdt*-*15* prevents animals from living long at low temperatures. We show that MDT-15 up-regulates *fat*-*7*, a fatty acid desaturase, at low temperatures, which increases the ratio of unsaturated to saturated fatty acids. We further demonstrate that maintaining this increased fatty acid ratio is essential for protein homeostasis and longevity at low temperatures. Thus, the homeostasis of lipid composition by MDT-15 appears to be a limiting factor for *C. elegans* proteostasis and longevity at low temperatures. Our findings highlight the crucial roles of fat regulation in maintaining normal organismal physiology under different environmental conditions.

## Introduction

Environmental temperature has a major influence on organismal physiology, including growth, metabolism, and aging. The body temperature of poikilothermic organisms, such as *Caenorhabditis elegans*, is subject to changes in environmental temperatures; these organisms live long at low ambient temperatures but short at high temperatures (Conti, 2008; Jeong et al., 2012). In addition, mice with reduced body temperature live long (Conti et al., 2006), suggesting potentially conserved mechanisms across diverse species. Several genetic factors have been identified to modulate temperature-dependent lifespan changes in *C. elegans* (Chen et al., 2016; Horikawa et al., 2015; Lee and Kenyon, 2009; Xiao et al., 2013). However, the metabolic processes underlying this phenomenon remain poorly understood.

The relative proportion of unsaturated fatty acids (UFAs) and saturated fatty acids (SFAs) is homeostatically regulated in various organisms (Holthuis and Menon, 2014). In *C. elegans*, the UFA/SFA ratio is inversely proportional to the environmental temperature (Tanaka et al., 1996). This is consistent with the idea that more UFAs than SFAs are required for the maintenance of physical properties of lipids in biological membranes at low temperature (Holthuis and Menon, 2014). Changes in the UFA/SFA ratio is also important for *C. elegans* to adjust its growth to high or low temperatures (Ma et al., 2015; Svensk et al., 2013). (Taubert et al., 2006; Van Gilst et al., 2005)However, the mechanisms by which the UFA/SFA ratio regulates the lifespan of organisms at different environmental temperatures remain largely unknown.

Here, we showed that MDT-15/Mediator 15, a subunit of the Mediator complex for RNA polymerase II-regulated transcription, has differential effects on lifespan at different environmental temperatures. We found that *mdt*-*15* mutations prevented worms from living long at low temperatures. We showed that *mdt*-*15* was required for expressing *fat*-*7*, a fatty acid desaturase crucial for increasing the UFA/SFA ratios at low temperatures. Furthermore, we demonstrated that a homeostatic increase in the UFA/SFA ratios at low temperatures was critical for longevity and chaperone-mediated proteostasis. These data suggest that MDT-15 is a limiting factor in the low temperature-mediated longevity of *C. elegans*, and it exerts this effect via the maintenance of the physical properties of lipids and protein homeostasis.

## Results

### *mdt*-*15* is required for longevity and organismal fitness at low temperatures

MDT-15/Mediator 15 is a large subunit of the Mediator complex that regulates diverse physiological processes, including fat metabolism, stress resistance, and lifespan (Allen and Taatjes, 2015). While performing experiments for our previous reports (Artan et al., 2016; Lee et al., 2015), we found that reduction of function *mdt*-*15* mutations greatly suppressed the long lifespan of *C. elegans* at a low temperature (15°C) (Fig. 1A; Supplementary Fig. S1A). In contrast, *mdt*-*15(*–) mutations had a marginal effect on the lifespan at a high temperature (25°C) (Fig. 1A; Supplementary Fig. S1A). To further confirm this result, we employed an auxin-inducible degron system using CRISPR/Cas9 knock in (Zhang et al., 2015), and generated *mdt*-*15::degron::EmGFP* strains (Fig. 1B). We found that auxin-induced depletion of MDT-15 (Fig. 1C; Supplementary Fig. S1B, C) substantially suppressed longevity at 15°C (Fig. 1D; Supplementary Fig. S1D). Next, we examined the effects of an *mdt*-*15* gain of function (*gof*) mutation (Svensk et al., 2013), which we generated in a wild-type background by using CRISPR, on lifespan at different temperatures. We found that the *mdt-15(gof*) mutations did not affect the lifespan at 25°C or 15°C (Fig. 1E). Together, these data indicate that MDT-15 is required but not sufficient for increasing the adult lifespan at low temperatures.

**Figure 1.**
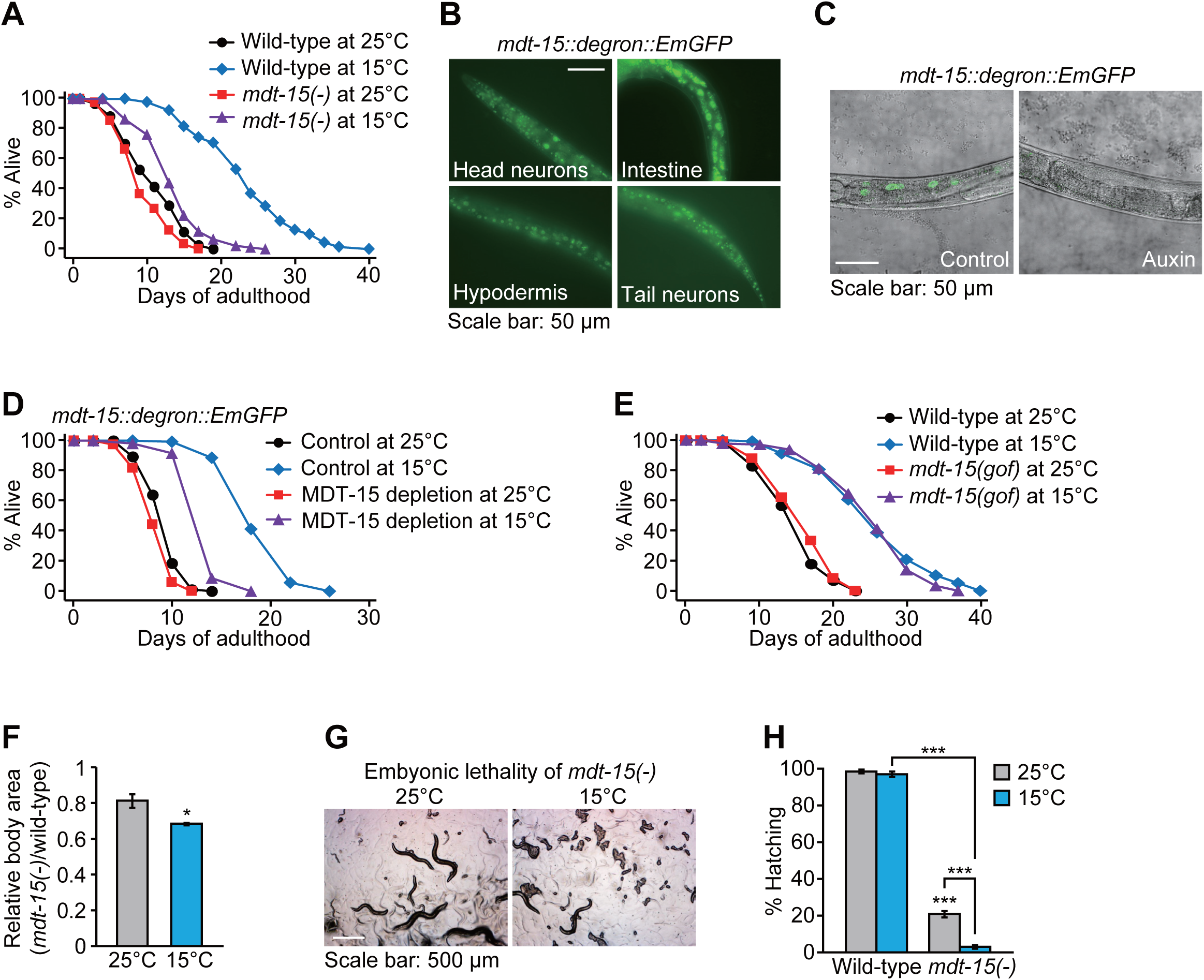
MDT-15 limits the body size and longevity at low temperatures. (**A**) Lifespan of wild-type and *mdt*-*15(tm2182)* [*mdt*-*15(–)*] *C. elegans* at 25°C and 15°C. Worms were cultured at indicated temperature from hatching. The lifespan assays in panel **A** were performed without 5-fluoro-2’-deoxyuridine (FUdR), which prevents progeny from hatching. (**B**) Expression of *mdt*-*15::degron::EmGFP* generated by CRISPR/Cas9 knock-in in the nuclei of many tissues, including neurons, intestine and hypodermis at 20°C. (**C**) Depletion of *MDT*-*15::degron::EmGFP* by treatment with auxin at 15°C was captured by using confocal microscopy. The worms carry ubiquitously expressed TIR1 (*eft*-*3p::TIR1::mRuby)* for the auxin-inducible protein degradation. (**D**) MDT-15 depletion substantially decreased longevity at 15°C. Control and MDT-15 depletion indicate solvent (ethanol) and auxin treatments, respectively. (**E**) Lifespan curves of wild-type and *mdt*-*15(yh8)* [*mdt*-*15(gof)*] mutant animals at 25°C and 15°C. (**F**) The relative ratio of body areas of *mdt*-*15(*–) and wild-type animals at 25°C and 15°C. The relative ratios were calculated from the same data set shown in Supplementary Fig. S1E. Bars indicate the averages of the ratios from three independent experiments. Eight worms were analyzed in each experimental set. Error bars represent standard error of the mean (SEM) (two-tailed Student’s t-tests, ^∗^*p*<0.05). (**G**) Images represent unhatched progeny in *mdt*-*15(*–) at 15°C. (**H**) Percent wild-type and *mdt*-*15(*–) mutant larvae that hatched at 25°C and 15°C. The hatching defect caused by *mdt*-*15(*–) was also measured at 16°C (Supplementary Fig. S1F). Error bars represent SEM (two-tailed Student’s t-tests, ^∗∗∗^*p*<0.001, n=248 for wild-type at 25°C, n=250 for wild-type at 15°C, n=644 for *mdt*-*15(*–) at 25°C, and n=1524 for *mdt*-*15(*–) at 15°C from five independent experiments). We were able to maintain *mdt*-*15(*–) worms at low temperatures, as the embryonic lethality was not 100%. See Supplementary Table 1 for statistical analysis and additional repeats of the lifespan assays.

Next, we determined whether *mdt*-*15* is important for maintaining the overall fitness, including the growth and reproduction of *C. elegans* at different temperatures. *mdt*-*15(*–) mutants exhibited a greater impact on body size at low temperatures than at high temperatures (Fig. 1F; Supplementary Fig. S1E). In addition, the defects in reproduction caused by *mdt*-*15(*–) mutations were more pronounced at a low temperature (15°C) than at a high temperature (25°C) (Fig. 1G, H; Supplementary Fig. S1F). Together, these data suggest that MDT-15 plays a role in longevity and fitness more substantially at low temperatures than at high temperatures.

### MDT-15 regulates the proper expression of fatty acid desaturases at low temperatures

Next, we sought to identify genes whose expression was affected by MDT-15 at different temperatures. Our RNA seq. analysis indicates that 79 genes were up-regulated and 253 genes were down-regulated at 15°C in an MDT-15-dependent manner (fold change > 1.5 and *p* values < 0.05; Fig. 2A–C; Supplementary Fig. S2A–F). Gene ontology (GO) analysis showed that GO terms involved in diverse metabolic processes, including aminoglycan, fatty acid, and carboxylic acid metabolism, were enriched in MDT-15-dependent up-regulated genes at 15°C (Fig. 2D). GO terms, including chromatin assembly, aminoglycan catabolic process, response to heat, immune system process, and response to unfolded protein, were overrepresented among the MDT-15-dependent down-regulated genes at 15°C (Fig. 2E).

**Figure 2.**
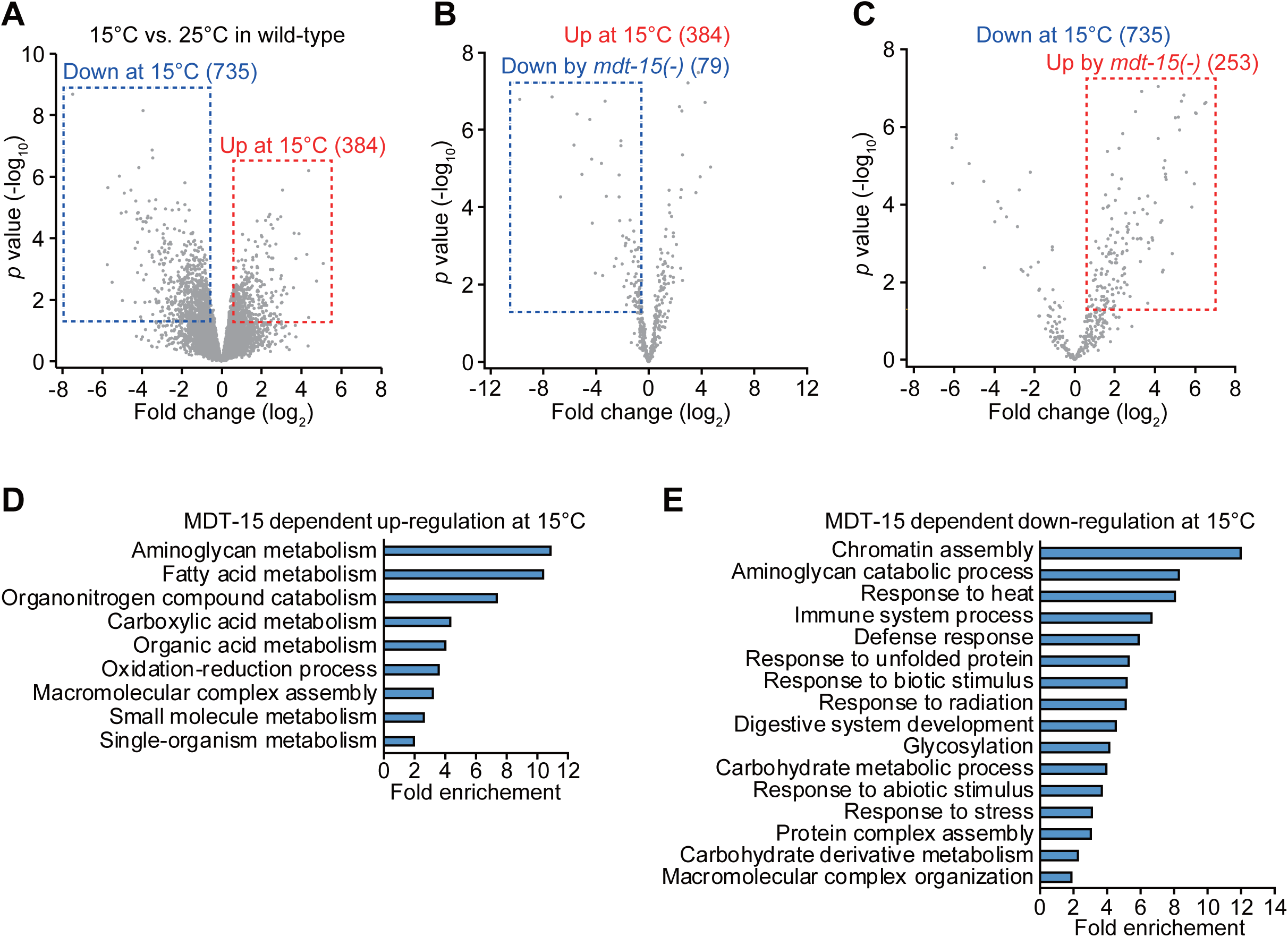
Genes that are differentially regulated by MDT-15 at different temperatures. (**AC**) Volcano plots represent genes up- and down-regulated in wild-type at 15°C (**A**), and genes up-regulated at 15°C (**B**) and down-regulated at 15°C (**C**) in an MDT-15 dependent manner. (**D, E**) Gene ontology (GO) terms enriched among the genes up-regulated at 15°C (**D**) and down-regulated at 15 °C (**E**) in an MDT-15-dependent manner.

Based on our GO analysis showing “metabolism” terms enriched among genes up-regulated at 15°C in an MDT-15-dependent manner, we further analyzed our RNA seq. data with a focus on genes located in carbohydrate and lipid metabolic pathways (Fig. 3A). We found that genes that directly affect lipid metabolism, including fatty acid synthesis, lipolysis, lipid transport, and fatty acid β-oxidation, tended to be more up- or down-regulated by changes in temperature and *mdt*-*15(*–) mutation than those involved in glycolysis and the Krebs cycle (Fig. 3A). In addition, the expression of many fatty acid desaturases and elongases involved in fatty acid biosynthesis was down-regulated in *mdt*-*15(*–) mutants, except *elo*-*8* (Fig. 3A). In contrast, the expression of genes involved in lipolysis, lipid transport, and fatty acid β-oxidation did not show a consistent pattern at different temperatures or by *mdt*-*15(*–) mutations (Fig. 3A). These data suggest that MDT-15 up-regulates fatty acid biosynthesis and desaturation at low temperatures.

**Figure 3.**
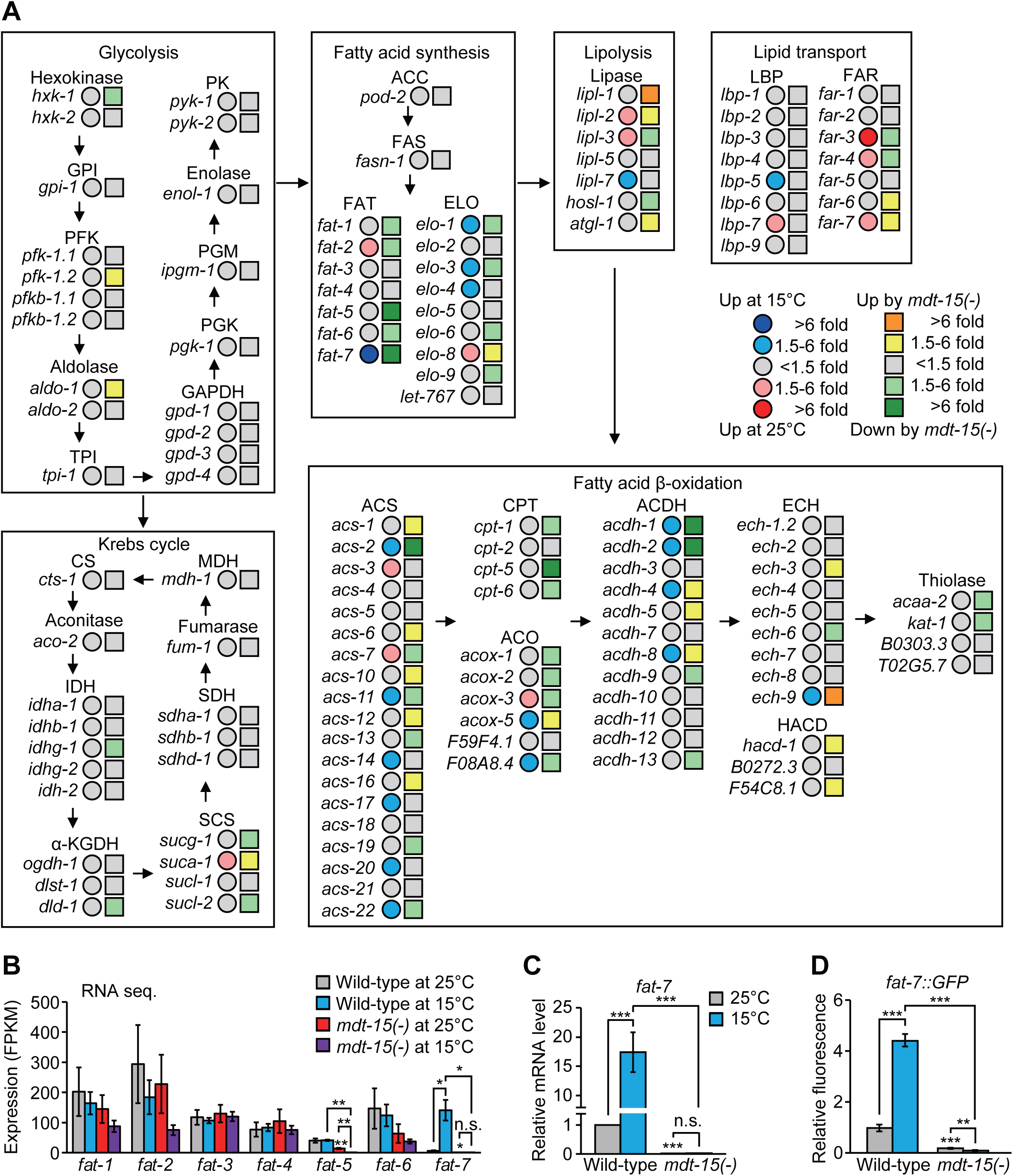
The effects of temperature changes and *mdt*-*15(*–) mutations on the expression of genes involved in metabolism. (**A**) Simplified carbohydrate and lipid metabolic pathways. Changes in the expression of genes involved in carbohydrate and fatty acid metabolism are shown. Fold changes at 15°C and by *mdt*-*15(tm2182)* [*mdt*-*15(*–)] mutations are shown in circles and squares, respectively. GPI: glucose-6-phosphate isomerase; PFK: phospho-fructo-kinase; TPI: triosephosphate isomerase; GAPDH: glyceraldehyde 3-phosphate dehydrogenase; PGK: phosphoglycerate kinase; PGM: phosphoglycerate mutase; PK: pyruvate kinase; CS: citrate synthase; IDH: isocitrate dehydrogenase; α-KGDH: α-ketoglutarate dehydrogenase; SCS: succinyl-CoA synthetase; SDH: succinate dehydrogenase; MDH: malate dehydrogenase; ACC: acetyl-CoA carboxylase; FAS: fatty acid synthase; FAT: fatty acid desaturase; ELO: fatty acid elongase; LBP: lipid-binding protein; FAR: fatty acid- and retinol-binding protein; ACS: acyl-CoA synthetase; CPT: carnitine palmitoyl transferase; ACO: acyl-CoA oxidase; ACDH: acyl-CoA dehydrogenase; ECH: enoyl-CoA hydratase; HACD: hydroxyacyl-CoA dehydrogenase. (**B**) mRNA levels of seven fatty acid desaturases in wild-type and *mdt*-*15(*–) at 25°C and 15°C were analyzed from RNA seq. data. (**C**) mRNA levels of*fat*-*7* in wild-type and *mdt*-*15(*–) at 25°C and 15°C were measured by using quantitative RT-PCR (qRT-PCR) (N=9, biological replicates). (**D**) Quantified fluorescence intensity of*fat*-*7::GFP* in wild-type and *mdt*-*15(*–) animals at 25°C and 15°C. Representative images are shown in Supplementary Fig. taG. Error bars represent SEM (two-tailed Student’s t-tests, ^∗^*p*<0.05, ^∗∗^*p*<0.01, ^∗∗∗^*p*<0.001, n=21 for wild-type at 25°C, wild-type at 15°C, *mdt*-*15(*–) at 25°C, and n=20 for *mdt*-*15(*–) at 15°C from three independent experiments)

We noticed that the expression of *fat*-*7* was the most strongly associated with temperature and MDT-15 genetic background changes (Fig. 3A, B). Among all seven fatty acid desaturases, the expression level of *fat*-*7* was significantly higher at 15°C than at 25°C (Ma et al., 2015), and this was MDT-15 dependent (Fig. 3B). In contrast, the other six fatty acid desaturase genes did not display MDT-15-dependent changes in the expression at different temperatures (Fig. 3B). qRT-PCR data for *fat*-*2*, *fat*-*5*, *fat*-*6*, and *fat*-*7* mRNAs and *fat*-*7::GFP* transgenic animals were generally consistent with our RNA seq. data (Fig. 3C, D; Supplementary Fig. S2G–J). Together, these data suggest that MDT-15 specifically increases the expression of *fat*-*7* at low temperatures.

### *mdt*-*15* mutations decrease the UFA/SFA ratio

Fatty acid desaturases and MDT-15 are crucial for *de novo* fat synthesis (Grants et al., 2015). By using Oil red O staining, we showed that the overall fat levels were not changed at 15°C in wild-type animals, but were greatly reduced in *mdt*-*15(*–) mutants at 15°C (Fig. 4A, B). At 15°C, *mdt*-*15(*–) mutants also displayed a pale intestine phenotype (Fig. 4C), which correlates with low fat levels (McKay et al., 2003). These data suggest that MDT-15 is crucial for maintaining overall fat levels at low temperatures.

**Figure 4.**
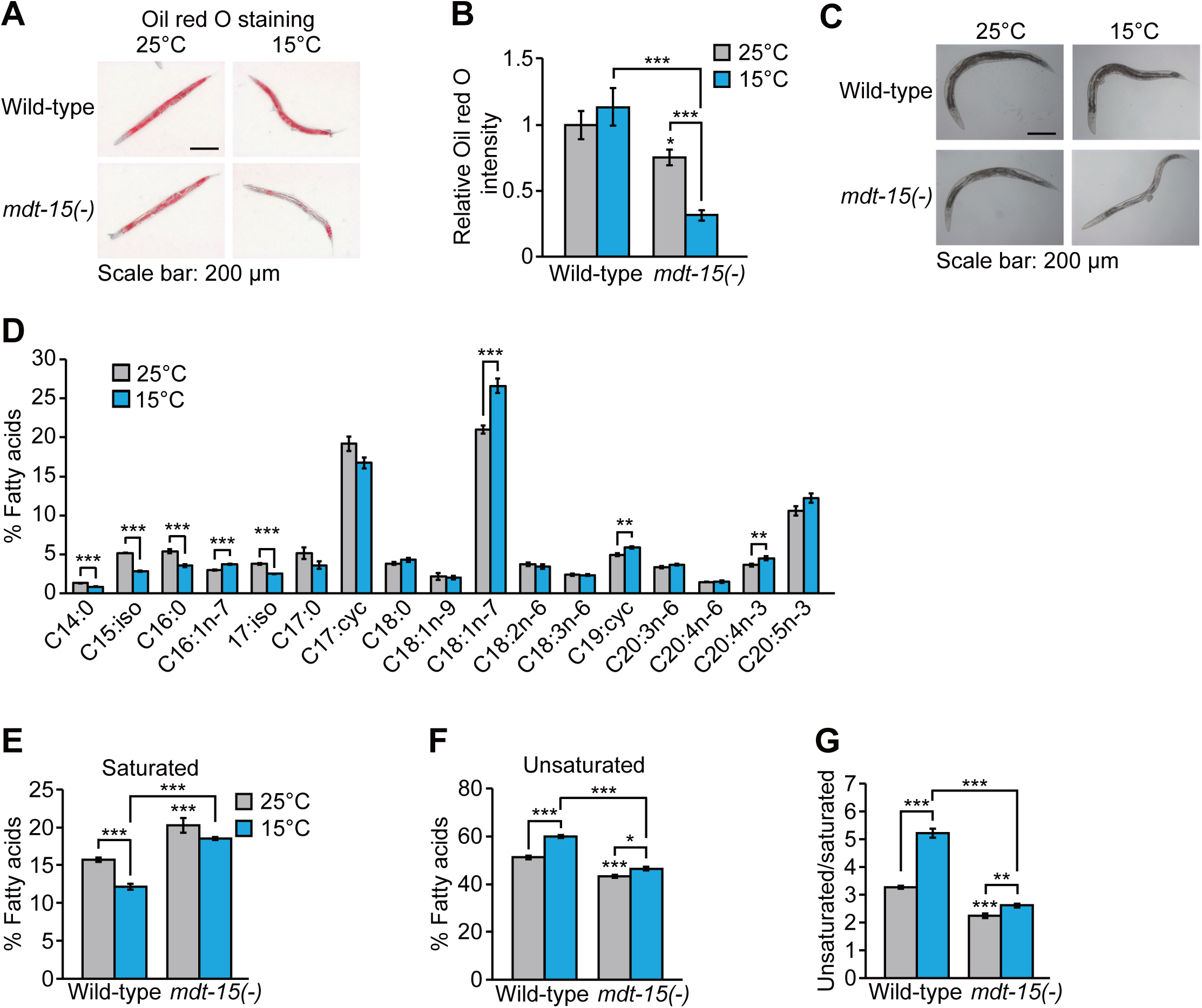
MDT-15 increases overall fat levels and UFA/SFA ratios at low temperatures. (**A**) Oil red O signals represent overall fat levels in wild-type and *mdt*-*15(tm2182)* [*mdt*-*15(*–)] mutant animals at 25°C and 15°C. (**B**) Quantification of the Oil red O staining in panel **A**. Error bars represent SEM (two-tailed Student’s t-tests, ^∗∗∗^*p*<0.001, n=20 for wild-type at 25°C, n=23 for wild-type at 15°C, n=20 for *mdt*-*15(*–) at 25°C, and n=24 for *mdt*-*15(*–) at 15°C from three independent experiments). (**C**) Bright field images of wild-type and *mdt*-*15(*–) mutants at 25°C and 15°C. *mdt*-*15(*–) mutant animals displayed pale intestines at 15°C. The images were taken when the worms reached day 1 adult stages. (**D**) Total fatty acid composition in wild-type animals at 25°C and 15°C was analyzed by using gas chromatography/mass spectrometry (GC/MS) (N=5, biological replicates). Fractions of each fatty acid in the total fatty acids (mole/mole) were calculated. (**E, F**) Fractions of saturated fatty acids (**E**) and unsaturated fatty acids (**F**) in wild-type and *mdt*-*15(*–) mutant animals at 25°C and 15°C. (**G**) The ratios of unsaturated fatty acids/saturated fatty acids that were calculated from the panels **E** and **F**. Error bars represent SEM (two-tailed Student’s t-tests, ^∗^*p*<0.05, ^∗∗^*p*<0.01, ^∗∗∗^*p*<0.001, n=5).

We then examined the effects of low temperature and *mdt*-*15(*–) mutations on the relative proportion of SFAs and UFAs because fatty acid desaturases increase the unsaturated status of fatty acids. The levels of SFAs, including C14:0 and C16:0, were reduced at 15°C (Fig. 4D); this was consistent with the findings of a previous report (Tanaka et al., 1996). In contrast, the levels of several UFAs, including C16:1n-7, C18:1n-7, and C20:4n-3, were increased at 15°C (Fig. 4D). The overall levels of SFAs were decreased, whereas those of UFAs were increased in the wild-type at 15°C (Fig. 4E, F), and this led to significant increases in the UFA/SFA ratio (Fig. 4G). Notably, *mdt*-*15(*–) mutations reduced the effect of low temperature on changes in the SFA and UFA levels (Fig. 4E, F). This resulted in a reduced UFA/SFA ratio in *mdt*-*15(*–) mutants, which was more pronounced at 15°C than at 25°C (Fig. 4G). These data suggest that MDT-15 is crucial for increasing the UFA/SFA ratio at low temperatures.

### Low UFA/SFA ratio suppresses longevity at low temperatures

Having established that *mdt*-*15* is crucial for maintaining overall fat levels and increasing the UFA/SFA ratio at low temperature, we sought to determine which of these two effects was critical for low temperature-induced longevity. Several lines of evidence based on genetic and dietary interventions suggest that decreasing the UFA/SFA ratio results in a short lifespan at low temperatures rather than a reduction in overall fat levels. First, *fat*-*6(*–); *fat*-*7(*–) double mutations, which increase SFA levels and decrease overall fat levels (Brock et al., 2007), significantly and specifically shortened lifespan at 15°C but not at 25°C (Fig. 5A). Second, mutations in the *paqr*-*2*/adiponectin receptor, which decrease the expression of *fat*-*7* and increase the levels of SFA (Svensk et al., 2013), largely suppressed longevity at 15°C (Fig. 5B). Third, *nhr*-*49* mutations, which decreased *fat*-*7* mRNA levels and the UFA/SFA ratio while increasing overall fat levels (Van Gilst et al., 2005) (Supplementary Fig. S3A-C), significantly shortened longevity at 15°C (Fig. 5C). The lifespan of *nhr*-*49*; *mdt*-*15* double mutants at low temperature was similar to that of the single mutants (Fig. 5D), suggesting that MDT-15 and NHR-49 act together to increase the longevity at low temperature. Fourth, glucose-enriched diets, which decrease the UFA/SFA ratio while increasing the overall fat levels (Devkota et al., 2017; Lee et al., 2015; Pang et al., 2014), also substantially shortened longevity at 15°C (Fig. 5E, F) but not at 25°C (Fig. 5E, F) (Brokate-Llanos et al., 2014; Lee et al., 2015). Taken together, these data suggest that increasing the UFA/SFA ratio, rather than maintaining the overall fat levels, is essential for the longevity of *C. elegans* at low temperatures.

**Figure 5.**
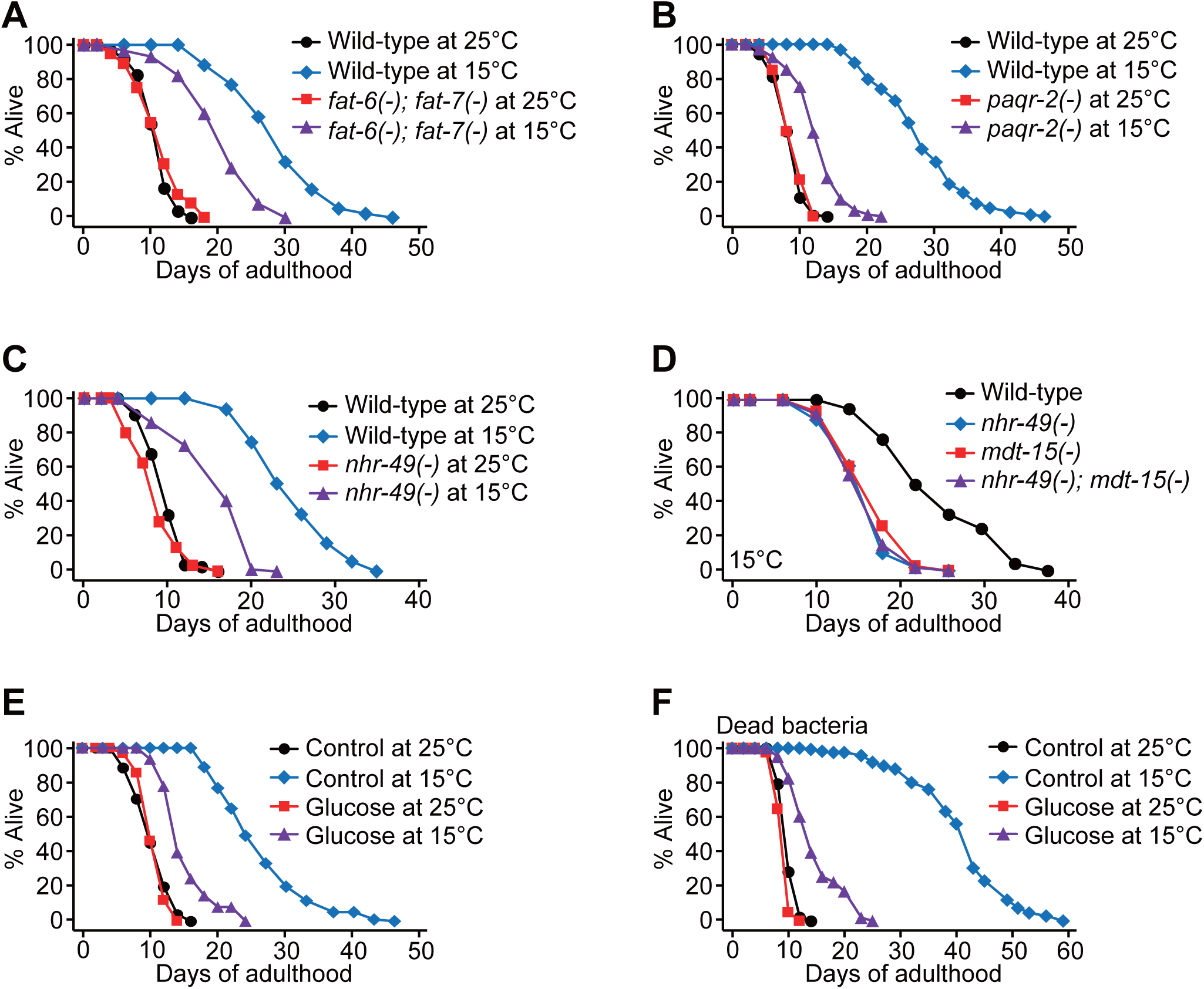
Maintenance of UFA/SFA ratios permits long lifespan at low temperatures. (**A**) Lifespan curves of wild-type and *fat*-*6(tm331)*; *fat*-*7(wa36)* [*fat*-*6(*–); *fat*-*7(*–)] mutants at 15°C and 25°C. Wild-type and*fat*-*6(*–); *fat*-*7(*–) mutants were cultured at 20°C until reaching L4 larval stages and then shifted to 15°C or 25°C. *mdt*-*15(tm2182)* [*mdt*-*15(*–)] mutations also substantially suppressed longevity at 15°C, when the worms were shifted from 20°C to 15°C at L4 larval stages (Supplementary Fig. S1A). (**B**) Lifespan curves of wild-type and *paqr*-*2(tm3410)* [*paqr*-*2(*–)] animals at 25°C and 15°C. Wild-type and*paqr*-*2(*–) worms were shifted from 20°C to indicated temperatures at L4 stages. (**C**) Lifespan curves of wild-type and *nhr*-*49(gk405)* [*nhr*-*49(*–)] animals at 25°C and 15°C. (**D**) *nhr*-*49(*–) mutations did not further shorten the lifespan of *mdt*-*15(*–) worms at 15°C. Genetic inhibition of *skn*-*1* or *sbp*-*1*, which encodes a transcription factor that interacts with MDT-15 (Goh et al., 2014; Pang et al., 2014; Yang et al., 2006), had a small or no effect on the longevity at low temperatures (Supplementary Fig. S3D, E). (**E**) Glucose-enriched diets (2%) substantially suppressed the longevity of wild-type animals at 15°C. (**F**) Lifespan curves of worms fed with dead bacteria supplemented with 2% glucose at 25°C and 15°C. These data indicate that the shortened lifespan upon glucose was not caused by changes in bacterial metabolism on glucose-rich media at different temperatures. The worms were shifted from 20°C to 25°C or 15°C at L4 stages. See Supplementary Table 1 for statistical analysis and additional repeats of the lifespan assays.

### Disruption of MDT-15-mediated fatty acid desaturation decreases proteostasis at low temperature

How does fatty acid composition affect longevity at low temperatures? In our RNA seq. analysis, we noticed that the GO terms “response to heat” and “response to unfolded protein” were enriched among genes down-regulated by low temperature in an MDT-15-dependent manner (Fig. 2E). We investigated the functional significance of the enrichment of these GO terms by focusing on our research at low temperature because protein folding and homeostasis are crucial for healthy aging and longevity (Higuchi-Sanabria et al., 2018). We found that reduced expression of cytosolic chaperones, such as *hsp*-*16.1*, *hsp*-*16.11*, *hsp*-*16.49*, *hsp*-*16.48*, *hsp*-*16.41*, *hsp*-*16.2*, *F44E5.4 (Hsp70)*, and *F44E5.5 (Hsp70)*, at 15°C was highly elevated in *mdt*-*15(*–) mutants (Fig. 6A). We confirmed this by measuring *hsp*-*16.1/11* and *F44E5.4/5* mRNA levels using *hsp*-*16.1::GFP* reporter and qRT-PCR (Fig. 6B-D). In contrast to the cytosolic chaperones, low temperatures or *mdt*-*15(*–) mutations had a small or no effect on the expression of endoplasmic reticulum or mitochondrial chaperones (Fig. 6E-G). Additionally, *paqr*-*2(*–) mutations induced the expression of *hsp*-*16.1/11* at 15°C (Supplementary Fig. S4). These data suggest that a reduction in the UFA/SFA ratio induces the expression of cytosolic chaperones at low temperatures.

**Figure 6.**
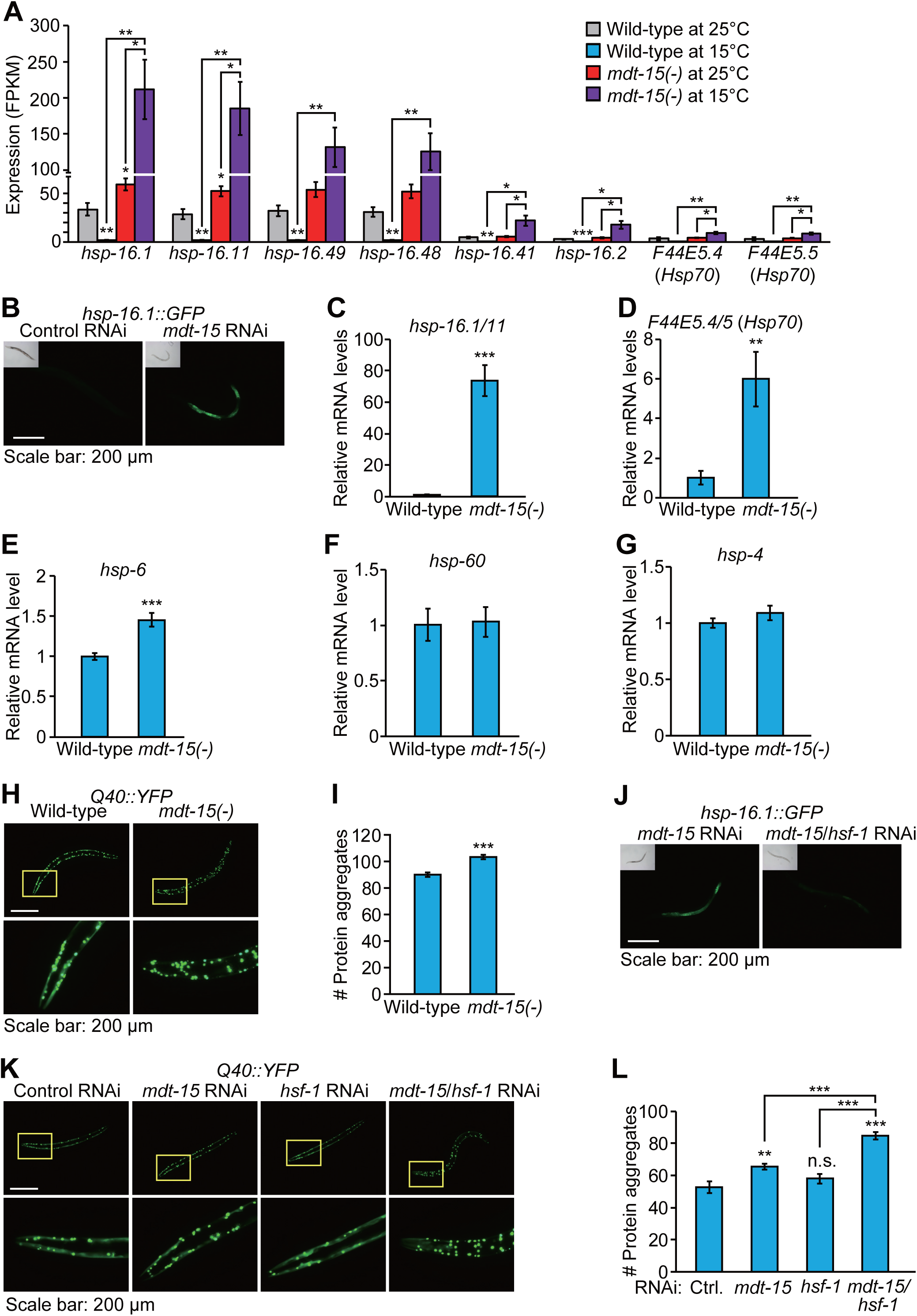
MDT-15 is required for maintaining proteostasis at low temperatures. (**A**) mRNA levels of selected cytosolic chaperones, *hsp*-*16.1*, *hsp*-*16.11*, *hsp*-*16.49*, *hsp*-*16.48*, *hsp*-*16.41*, *hsp*-*16.2*, *F44E5*.*4*, and *F44E5.5* in wild-type and *mdt*-*15(*–) worms at 25°C and 15°C were measured by using RNA seq (N=3, biological replicates). Note that the mRNA levels of *hsp*-*16.1* and *hsp*-*16.11*, those of *hsp*-*16.48* and *hsp*-*16.49*, and those of *F44E5.4* and *F44R5.5* may not be completely distinguishable, because the sequences of these pairs of transcripts are very similar. The induction of some of these chaperone genes by inhibition of *mdt*-*15* was also observed at 20°C in a previous report (Taubert et al., 2008). (**B**) Images of *hsp*-*16.1::GFP* transgenic worms treated with control RNAi or *mdt*-*15* RNAi at 15°C. (**C**) mRNA levels of *hsp*-*16.1/11* were measured by using qRT-PCR in wild-type and *mdt*-*15(tm2182)* [*mdt*-*15(*–)] animals at 15°C (N=7, biological replicates). (**D**) The expression of *F44E5.4/5* at 15°C was determined by using qRT-PCR (N=6, biological replicates). (**E-G**) qRT-PCR data represent mRNA levels of *hsp*-*6* (N=6, biological replicates) (**E**), *hsp*-*60* (N=6, biological replicates) (**F**), and *hsp*-*4* (N=5, biological replicates) (**G**) in wild-type and *mdt*-*15(*–) worms at 15°C. (**H**) Fluorescence images of Q40::YFP transgenic worms in wild-type and *mdt*-*15(*–) backgrounds at 15°C. The images were captures at L4 larval stages. (**I**) Quantification of the Q40::YFP aggregates shown in panel **H** (n =30 for all conditions from three independent experiments). (**J**) Fluorescence images of *hsp*-*16.1::GFP* treated with *mdt*-*15* RNAi or *mdt*-*15/hsf-1* double RNAi at 15°C. (**K**) Images of Q40::YFP transgenic worms treated with *mdt*-*15* RNAi, *hsf*-*1* RNAi, or *mdt*-*15/hsf*-*1* double RNAi at 15°C. The images were taken at L4 stages. (**L**) Quantification of the Q40::YFP aggregates shown in panel **K** (n=29 for control RNAi, and n=28 for *mdt*-*15* RNAi, *hsf*-*1* RNAi, and *mdt*-*15/hsf*-*1* RNAi from three independent experiments). Note that the number of Q40::YFP aggregation in control RNAi condition was smaller than wild-type condition in panel **I**. We speculate that different food types (OP50 vs. HT115) affect Q40 aggregation. Error bars represent SEM (two-tailed Student’s t-tests, ^∗^*p*<0.05, ^∗∗^*p*<0.01, ^∗∗∗^*p*<0.001, n.s.: not significant).

Next, we determined how the genetic inhibition of *mdt*-*15* increased the expression of cytosolic chaperones at low temperatures. We hypothesized that the inhibition of *mdt*-*15* at low temperature increased proteotoxicity, which in turn induced chaperone gene expression as an adaptive response. Consistent with our hypothesis, *mdt*-*15* mutations significantly increased the number of Q40::YFP puncta (Fig. 6H, I), a poly-glutamine proteotoxicity model in *C. elegans* (Morley et al., 2002). HSF-1/heat shock factor 1 is a major transcription factor that regulates the expression of cytosolic chaperones (Higuchi-Sanabria et al., 2018); *hsf*-*1* knockdown suppressed the induction of *hsp*-*16.1::GFP* by *mdt*-*15* RNAi at 15°C (Fig. 6J). Importantly, *hsf*-*1* knock down exacerbated Q40::YFP aggregation in *mdt*-*15* RNAi-treated worms but did not significantly affect the number of the aggregates in control worms at 15°C (Fig. 6K, L). These data imply that HSF-1-mediated chaperone induction is an adaptive response to increased proteotoxicity caused by the inhibition of *mdt*-*15* at 15°C. Altogether, our data suggest that the homeostatic regulation of lipid composition regulated by MDT-15 is crucial for maintaining proteostasis and longevity at low temperatures.

## Discussion

### UFA/SFA ratio limits longevity at low temperatures

In this study, we showed that *mdt*-*15* was required for the longevity of *C. elegans* at low temperatures. When worms were cultured at a low temperature (15°C), MDT-15 increased the UFA/SFA ratio by permitting the expression of several fatty acid desaturases, including *fat*-*7.* This homeostatic regulation of lipid metabolism by MDT-15 was essential for the longevity of *C. elegans* at low temperatures. Consistent with this possibility, *mdt*-*15(*–) mutations reduced the UFA/SFA ratio and suppressed the low temperature-induced longevity. Similar to *mdt*-*15(*–) mutations, *fat*-*6(*–); *fat*-*7(*–), *paqr*-*2(*–), and *nhr*-*49(*–) mutations and glucose-enriched diets suppressed longevity at low temperatures. The reduction in the UFA/SFA ratio at low temperatures led to proteotoxicity, resulting in shortened lifespan. Previous studies have demonstrated that modulating the UFA/SFA ratio is crucial for larval development or resistance against environmental stress caused by extreme fluctuations in temperature (Brock et al., 2007; Ma et al., 2015; Murray et al., 2007; Svensk et al., 2013). Our data suggest that the homeostatic regulation of the UFA/SFA ratio is essential for the maintenance of proteostasis and longevity in adult *C. elegans* at low temperatures.

### MDT-15 regulates lifespan in response to changes in temperature and diet

*C. elegans* MDT-15 is a transcriptional co-regulator involved in the physiological response to ingested materials, including food, toxins, pathogens, and stressors (Goh et al., 2014; Pukkila-Worley et al., 2014; Schleit et al., 2011; Taubert et al., 2008; Taubert et al., 2006; Yang et al., 2006). Previously, we showed that MDT-15-regulated lipid metabolism is crucial for maintaining normal lifespan in glucose-enriched nutrient conditions (Lee et al., 2015). In this study, we demonstrated that MDT-15 was important for longevity at low ambient temperatures. Together, these findings suggest that MDT-15 plays a key role in adapting to changes in environmental temperatures and to various ingested materials. How MDT-15 differentially incorporates various inputs to exert proper cellular and physiological responses currently remains elusive. Because MDT-15 is a subunit of the Mediator complex that interacts with diverse transcription factors (Allen and Taatjes, 2015), it is possible that MDT-15 binds to these transcription factors differently under different conditions.

### Some specific transcription factors acting with MDT-15 may regulate the temperature-dependent expression of *fat*-*7*

Among the seven *C. elegans* fatty acid desaturases, the expression of *fat*-*7* was down-regulated at high temperature, instead of being up-regulated at low temperature, in an MDT-dependent manner (Fig. 3B). This suggests that the activity of some transcription factors, including MDT-15, NHR-49, SBP-1, DAF-16/FOXO, and HSF-1, that induce *fat*-*7* (Brunquell et al., 2016; Murphy et al., 2003; Taubert et al., 2006; Yang et al., 2006) is reduced at high temperatures. The effect of temperature on the activity of these transcription factors has not been investigated, except for the heat-activated HSF-1. However, HSF-1 positively regulates the expression of *fat*-*7* at high temperatures (Brunquell et al., 2016), and this is in contrast to the data shown in this study. A recent study has shown that acyl-CoA dehydrogenase, ACDH-11, whose expression is up-regulated at high temperatures, negatively regulates *fat*-*7* via NHR-49 (Ma et al., 2015), which acts with MDT-15 for target gene expression (Taubert et al., 2006). Thus, NHR-49 with MDT-15 may regulate the temperature-dependent expression of *fat*-*7.*

### MDT-15-regulated membrane fluidity may influence lifespan

The UFA/SFA ratio is crucial for maintaining the membrane fluidity for adaptation to different temperatures (Holthuis and Menon, 2014). In *C. elegans*, the rigid membrane leads to developmental defects at low temperatures (Svensk et al., 2016) and hyper-fluidic membrane decreases resistance against high temperatures (Ma et al., 2015). Here, we showed that *mdt*-*15(*–) mutations decreased the UFA/SFA ratio at low temperature and suppressed the longevity. We also found that genetic interventions and glucose-enriched diets, which decrease the UFA/SFA ratio, shortened lifespan specifically at low temperatures. Thus, a reduction in the UFA/SFA ratio at low temperatures may increase the rigidity of membranes to a harmful level, which in turn may shorten lifespan. A recent study has demonstrated that feeding worms with glucose or glycolysis metabolites increases their membrane rigidity by altering bacterial metabolism (Devkota et al., 2017). However, we showed that feeding *C. elegans* with a glucose-enriched diet along with dead bacteria also shortened lifespan substantially at low temperature (Fig. 5F). Therefore, glucose-rich diets appear to directly shorten the lifespan of *C. elegans* at low temperatures independently of bacterial metabolism.

### MDT-15 regulates proteostasis independently of mitochondrial stress

Mitochondrial stress, or a mild disruption of mitochondrial functions, up-regulates the expression of cytosolic chaperones (Kim et al., 2016; Labbadia et al., 2017). Therefore, it is possible that the induction of cytosolic heat shock proteins in *mdt*-*15(*–) mutants observed in this study implicates mitochondrial stress. However, several lines of evidence are against this possibility. First, we found that *mdt*-*15(*–) mutations had a small or no effect on the induction of mitochondrial stress response genes. Second, unlike the increase in fat levels required for the induction of cytosolic chaperones under mitochondrial stress (Kim et al., 2016), we showed that *mdt*-*15(*–) mutations reduced overall fat levels while increasing cytosolic chaperone levels. Third, different from mitochondrial perturbations that improve proteostasis (Kim et al., 2016; Labbadia et al., 2017), we found that the inhibition of *mdt*-*15* disrupted proteostasis. Overall, our data suggest that the induction of cytosolic chaperones by *mdt*-*15(*–) mutations is a response to increased proteotoxicity and is distinct from mitochondrial stress.

### Low UFA/SFA ratios are associated with various human diseases

Recent reports indicate that supplementing with certain types of UFAs is sufficient to extend the lifespan of *C. elegans* (Han et al., 2017; O’Rourke et al., 2013). Together with our current report, these findings suggest the beneficial effects of UFAs on healthy aging. Interestingly, low UFA/SFA ratios are associated with many diseases such as Niemann-Pick disease, hypertension, heart disease, and Alzheimer’s disease (Chi and Gupta, 1998; Soderberg et al., 1991; Sztolsztener et al., 2012; Wang et al., 2003). Human fibroblasts that carry mutations in *NPC1*, the gene responsible for the Niemann-Pick type C disease, display high levels of SFAs and a reduced membrane fluidity (Sztolsztener et al., 2012). Low UFA/SFA ratios also positively correlate with heart disease incidences in humans (Wang et al., 2003). Spontaneously hypertensive rats show lower UFA/SFA ratio in aorta and kidney than normotensive rats (Chi and Gupta, 1998). Patients of Alzheimer’s disease show very high levels of SFAs in their brains (Soderberg et al., 1991), and this is consistent with our data that show the occurrence of proteotoxicity because of low UFA/SFA ratios. These findings point toward the importance of low UFA/SFA ratios in the pathophysiology of the disease. Further research on MDT-15 and lipid metabolism using *C. elegans* may provide useful information that may eventually help the treatment of human diseases because functions of the Mediator complex are evolutionarily well conserved (Allen and Taatjes, 2015).

### Homeostatic regulation of lipid metabolism is crucial for longevity

Homeostatic regulation of proteins and DNA is important to prevent premature aging and promote longevity (Lopez-Otin et al., 2013). In addition, recent studies indicate that RNA quality control mediated by nonsense-mediated mRNA decay and accurate mRNA splicing contributes to the longevity in *C. elegans* (Heintz et al., 2017; Son and Lee, 2017; Son et al., 2017; Tabrez et al., 2017). Our current findings suggest that lipid homeostasis is also crucial for longevity at low temperatures, perhaps because lipids are very susceptible to changes in environmental temperatures. We further showed that the disruption of lipid composition at low temperatures increased proteotoxicity and shortened lifespan. Thus, the homeostatic regulation of lipids and proteins is most likely to be interconnected and tightly regulated for health and longevity.

## Materials and Methods

### Strains

All strains were maintained on nematode growth medium (NGM) agar plates seeded with *E. coli* (OP50) at 20°C. Following strains were used in this study. N2 wild-type, IJ235 *mdt*-*15(tm2182) III* obtained by outcrossing XA7702 (a gift from Stefan Taubert lab) four times to Lee lab N2, IJ1648 *ieSi57[eft*-*3p::TIR1::mRuby::unc*-*54 3 ‘UTR; cb-unc-119] II* outcrossed four times to Lee lab N2, IJ1651 *yh44[mdt*-*15::degron::EmGFP] III* outcrossed four times to Lee lab N2 after CRISPR/Cas9 editing, IJ1729 *ieSi57[eft*-*3p::TIR1::mRuby::unc*-*54 3 ‘UTR; cb*-*unc*-*119] II; yh44[mdt*-*15::degron::EmGFP]III* obtained by crossing IJ1648 and IJ1651, IJ1468 *mdt*-*15(yh8) III* outcrossed four times to Lee lab N2 after CRISPR/Cas9 editing, DMS303 *nIs590[fat*-*7p::fat*-*7::GFP] V.* IJ1742 *mdt*-*15(tm2182) III; nIs590[fat*-*7p::fat*-*7::GFP] V* obtained by crossing IJ235 and DMS303, IJ511 *fat*-*6(tm331) IV; fat*-*7(wa36) V* outcrossed seven times to Lee lab N2, IJ666 *paqr*-*2(tm3410) III* obtained by outcrossing QC121 (a gift from Marc Pilon lab) four times to Lee lab N2, CF2774 *nhr*-*49(gk405) I* outcrossed four times to N2, IJ360 *nhr*-*49(gk405) I; mdt*-*15(tm2182) III*, obtained by crossing CF2774 and IJ235. IJ1625 *skn*-*1(zj15)* outcrossed four times to Lee lab N2. IJ414 *Is[hsp*-*16.1p::hsp*-*16::gfp; rol*-*6D]* (a gift from Junho Lee lab). IJ184 *rmIs133[unc*-*54p::Q40::YFP]* outcrossed four times to Lee lab N2. IJ1740 *mdt*-*15(tm2182) III; rmIs133[unc*-*54p::Q40::YFP]* obtained by crossing IJ235 and IJ184.

### CRISPR/Cas9 genome editing

*mdt*-*15::degron::EmGFP* knock-in was generated as described previously with modifications (Zhang et al., 2015). The *degron::EmGFP* repair template with homology arms (37 bp for 5’ and 36 bp for 3’) was amplified by PCR using pLZ29 as a template (Zhang et al., 2015), and the PCR products were purified by using PCR purification kit (QIAGEN, Germany). Wild-type adult worms were injected with pRF4 (roller injection marker, 50 ng/μl), Cas9 protein (250 ng/μl, Integrated DNA Technologies, USA), tracrRNA (100 ng/μl, Integrated DNA Technologies, USA), *mdt*-*15* crRNA (56 ng/μl, ATAATCTTAACTTGTAAGTT), and the purified *degron::EmGFP* repair template (450 ng/μl). Individual worms were subsequently transferred to new plates, and roller and non-roller F1 worms from plates that contained many rollers were genotyped by using PCR to identify worms that contained knock-in mutations. The *mdt*-*15::degron::EmGFP* knock-in was confirmed by sequencing, and the strain was outcrossed four times to Lee lab N2 to remove potential background mutations. For generating *mdt*-*15(yh8*, *gof)* (Svensk et al., 2013) in wild-type background, CRISPR/Cas9 plasmids targeting *mdt*-*15* were generated by replacing *pha*-*1* sgRNA sequence in pJW1285 (Ward 2015) with *mdt*-*15* sgRNA sequence (TTTCTTGCCTGAGCTGATGT). Wild-type adults were injected with pRF4 (roller injection marker, 50 ng/μl), pIJ285 (CRISPR/Cas9 plasmid targeting *mdt*-*15*, 50 ng/μl), and repair template (ATCGAGCTCCTGTGCCTCCAGATCCACAACTAACATCAGCTCAGGCAAGAAATCCACCTGTTACCGTAGCA, 10 ng/μl), and individual worms were transferred to new plates. F1 roller worms were genotyped by PCR to identify the *mdt*-*15(gof)* mutation. The *mdt*-*15(gof)* mutation was confirmed by using sequencing, and the *mdt*-*15(gof)* mutant worms were outcrossed four times to wild-type worms to remove potential non-specific mutations.

### Lifespan assays

Lifespan assays were performed as described previously with some modifications (Lee et al., 2010). Synchronized worms were cultured on the OP50-seeded NGM plates at different temperatures, and then transferred to plates containing 10 μM 5-fluoro-2’-deoxyuridine (FUdR, SIGMA, USA) at young adult stage (day 1) to prevent their progeny from hatching. Worms were transferred to fresh plates every other day until they stopped producing progeny for lifespan assays without FUdR treatment. Auxin-inducible degron assays were performed as described (Zhang et al., 2015). Briefly, synchronized worms were cultured on OP50-seeded NGM plates at 25°C or 15°C until reaching day 1 adult stage, and then transferred to new OP50-seeded NGM plates containing 1 mM auxin (indole-3-acetic acid, ALFA AESAR, USA). Ethanol (solvent) was used as a control for auxin treatments. For temperature shift assays, worms were cultured at 20°C until reaching L4 stage, and subsequently shifted to 25°C or 15°C. When the worms reached young adult stage (day 1), the worms were transferred to plates containing FUdR. For glucose-enriched diet experiments, 2% glucose (Junsei Cheminal, Japan) was added to NGM as described previously (Lee et al., 2009). OP50 bacteria that were cultured overnight in liquid LB media were concentrated 20 times by centrifugation and seeded onto NGM plates containing 10 μg/ml kanamycin (SIGMA, USA) for dead bacteria assays as previously described (Lee et al., 2009). For RNAi experiments, bacteria that express double stranded RNA targeting a specific gene were cultured in liquid LB containing 50 μg/ml ampicillin (USB, USA) at 37°C overnight, and then seeded onto NGM containing 50 μg/ml ampicillin. The RNAi bacteria-seeded plates were incubated at 37°C overnight, and treated with 1 mM isopropyl β-D-1-thiogalactopyranoside (IPTG, GOLDBIO, USA) to induce double stranded RNA at room temperature. Deaths of worms were determined by no response upon gently touching with a platinum wire. Animals that crawled off the plates, ruptured, bagged, or burrowed were censored but included in the statistical analysis. Days of adulthood are presented in lifespan curves, and time zero in lifespan data indicates a time when worms reached adulthood.

### Body size assays

Wild-type and *mdt*-*15(tm2182)* worms were cultured on the OP50-seeded HG (high growth NGM) plates at 20°C, and synchronized by using a bleaching method (Stiernagle, 2006). The bleached eggs were kept in M9 buffer at 20°C overnight, and then hatched L1 worms were cultured on OP50-seeded NGM plates at 25°C or at 15°C. After becoming fully-grown adults (72 hours at 25°C and 144 hours at 15°C), the worms were placed on a 2% agarose pad and anesthetized by using 100 mM sodium azide (SIGMA, USA). Bright field images were captured using AxioCam HRc (ZEISS, Germany) camera attached to a Zeiss Axioscope A.1 microscope (ZEISS, Germany). ImageJ (Schneider et al., 2012) was used for the quantification of body areas.

### Reproduction assays

Wild-type and *mdt*-*15(tm2182)* worms were cultured at 25°C or 15°C at least two generations. Single L2- or L3-staged larvae were transferred onto new OP50-seeded NGM plates and kept at 25°C or 15°C. Twenty wild-type and thirty *mdt*-*15* mutant worms at each temperature were used for one experimental set, and the assays were repeated five times. When the worms were producing eggs (for 2-3 days at 25°C and 4-5 days at 15°C), the eggs were transferred onto new OP50-seeded NGM plates, and kept at 25°C or 15°C. When the hatched larvae reached L3-L4 stages, larva/egg ratios were calculated for embryonic lethality. For 25°C and 16°C reproduction experiments, worms were first grown at 20°C to gravid adult stage. Ten worms were then transferred to 25°C and 16°C, respectively, and allowed to lay eggs for 24 hours before being removed. The eggs remained at their respective temperatures for another 24 hours to allow them to hatch. The number of live progeny and dead eggs was then counted after another 24 hours.

### RNA seq. analysis

HG-cultured wild-type and *mdt*-*15(tm2182)* worms were synchronized by using a bleaching method (Stiernagle, 2006) and incubated in M9 buffer at 20°C for overnight. The L1 worms were transferred onto OP50-seeded NGM plates and incubated at 25°C or 15°C until reaching day 1 adult stage. The adult worms were harvested by washing twice with M9 buffer, and then frozen at −80°C. Total RNA of the worms was extracted by using RNAiso plus (TAKARA, Japan). Three independent biological repeats were used for analysis. cDNA library was prepared and cDNA sequencing was performed by using a HiSeq 4000 platform by MACROGEN (MACROGEN, South Korea). Paired-end reads were aligned to the *C. elegans* genome ce11, and analyzed by using HISAT2 (v.2.0.5), StringTie (v.1.3.3), and Ballgown (v.2.0.0) methods (Pertea et al., 2016). R packages limma (v.3.24.15) (Ritchie et al., 2015) and edgeR (v.3.10.5) (Robinson et al., 2010) were used for analyzing differentially expressed genes (DEG), and genes whose expression was not detected (counts per million (cpm) < 1) were excluded. Genes with significant changes in expression (fold change > 1.5 and *p* values < 0.05) were further analyzed. Heat maps were generated by using Cluster 3.0 (de Hoon et al., 2004) and Java Treeview (Saldanha, 2004). Venn diagrams were generated by using Venn Diagram Plotter (http://omics.pnl.gov/software/venn-diagram-plotter). Gene ontology assays were performed by using DAVID (Huang da et al., 2009). Changes in the expression of metabolic genes were displayed as previously shown with modifications (Steinbaugh et al., 2015; Yu et al., 2017). Genes that were excluded during the DEG analysis are not presented in the metabolic pathways.

### Quantitative RT-PCR

Quantitative RT-PCR was performed as described previously with modifications (Seo et al., 2013). Synchronized worms were cultured at different temperatures and were harvested at young adult stage (day 1) by washing twice with M9 buffer. *paqr*-*2(tm3410)* and control (wild-type) worms were cultured at 20°C until reaching L4 stages, and then shifted to 25°C or 15°C, for maintaining consistency with lifespan assays. Total RNA was isolated using RNAiso plus (TAKARA, Japan) and reverse transcription was performed using ImProm-II^™^ Reverse Transcriptase kit (PROMEGA, USA). Random primers (9-mers, Cosmogenetech, South Korea) were used for reverse transcription. Quantitative real time PCR was performed by using StepOne and StepOnePlus Real time PCR system (APPLIED BIOSYSTEMS, USA). Relative quantity of specific mRNA was analyzed by using comparative Ct methods described in the manufacturer’s manual, and the mRNA level of *ama*-*1*, which encodes an RNA polymerase II large subunit, was used for normalization. The average of two technical repeats was used for each biological dataset.

### Primers used for quantitative RT-PCR assays

*ama-1*-F: TGGAACTCTGGAGTCACACC

*ama-1*-R: CATCCTCCTTCATTGAACGG

*fat-2*-F: AGTTTCTGGAGTTGCATGCGCTATC

*fat-2*-R: CAGCTTCGTAGACCTCAATATCCTC

*fat-5*-F: GTTCCAGAGGAAGAACTACCTCCCC

*fat-5*-R: GGGTGAAGCAGTAACGGAAGAGGGC

*fat-6*-F: GCGCTGCTCACTATTTCGGATGG

*fat-6*-R: GTGGGAATGTGTGATGGAAGTTGTG

*fat-7*-F: CTGCACGTCGCCGCAGCCATTG

*fat-7*-R: GAGAGCAAATGAGAAGACGGCC

*hsp-16.1/11*-F: CTCATGAGAGATATGGCTCAG

*hsp-16.1/11*-R: CATTGTTAACAATCTCAGAAG

*F44E5.4/F44E5.5*-F: GAATGGAAAGGTTGAGATCCTC

*F44E5.4/F44E5.5*-R: CCAACCAATCTTTCCGTATCTG

*hsp-6*-F: CTATGGGCCCAAAAGGAAGAAACGTG

*hsp-6*-R: GGGAATACACTTTTCCTTGAGCCTC

*hsp-60*-F: CTATGGGCCCAAAAGGAAGAAACGTG

*hsp-60*-R: GGATTTCGCGACGGTGACTCCGTCC

*hsp-4*-F: TCAGAAACTTCGCCGTGAGGT

*hsp-4*-R: AGAGTGACTCGATCTCGATCT

### Oil red O staining

Oil Red O staining was performed as described previously with some modifications (O’Rourke et al., 2009). Synchronized worms were cultured on OP50-seeded NGM plates at 25°C or 15°C. Approximately 300 young adult (day 1) worms were harvested by washing twice with M9 buffer and fixed with 60% isopropanol for two minutes. Oil Red O solution (0.5% in isopropanol, SIGMA, USA) was diluted in double distilled water (ddH_2_O) to prepare 60% working solution. Precipitates of Oil red O were eliminated by filtering. The fixed worms were incubated in the 60% working solution overnight at 25°C. The stained worms were washed with M9 buffer, and subsequently were placed on a 2% agarose pad with M9 buffer containing 0.01% Triton X-100 (DAEJUNG, South Korea) using a micropipette. DIC images were captured using AxioCam HRc (ZEISS, Germany) camera attached to a Zeiss Axioscope A.1 microscope (ZEISS, Germany). ImageJ (Schneider et al., 2012) was used for the quantification of Oil Red O intensity. The backgrounds of the images were subtracted and the images were then converted to 8-bit grayscale images. Same thresholds were set in the same experimental sets for detecting the Oil red O signals. Areas that had higher intensities than the threshold in the whole body were measured, and the areas were normalized by body size.

### Fluorescence imaging

GFP transgenic worms were anesthetized by using 100 mM sodium azide, and then placed on a 2% agarose pad. Images of the worms were captured by using AxioCam HRc (ZEISS, Germany) camera attached to a Zeiss Axioscope A.1 microscope (ZEISS, Germany). ImageJ (Schneider et al., 2012) was used to quantify the fluorescence intensity and the background signals were subtracted. High resolution confocal images were captured by using the Nikon A1si/Ni-E upright confocal microscope with 60×, 1.4 NA oil-immersion objective lens in the Advanced Neural Imaging Center in Korea Brain Research Institute (KBRI). For double RNAi treatments, control and gene-specific RNAi bacteria were separately cultured in liquid LB containing 50 μg/ml ampicillin (USB, USA) at 37°C. OD590 was adjusted to 0.9, and same volumes of the bacterial culture were mixed as described previously (Rea et al., 2007). Gene-specific RNAi bacteria were mixed with control RNAi bacteria for single RNAi treatments in the same experimental sets for comparison.

### Fatty acid composition assays

Lipid extraction was performed as described previously with some modifications (Deline et al., 2013). Wild-type and *mdt*-*15(tm2182)* worms were synchronized by using a bleaching method (Stiernagle, 2006) and incubated in M9 solution for overnight at 20°C. Hatched L1 worms were then cultured on OP50-seeded NGM plates at 25°C or 15°C. Approximately 1000-1500 day 1 adult worms were harvested and washed three times with ddH_2_O. The harvested worms were frozen in liquid nitrogen and stored at −80°C until use. Fatty acid methyl esters (FAMEs) were prepared by adding 2 ml of sulfuric acid (2.5%)/methanol solution to the worm samples, followed by heating at 70°C for one hour. The FAMEs were extracted by using 3 ml ddH_2_O and 3 ml of hexane. The hexane layers were evaporated and analyzed by using GC/MS (GCMS-QP2010, Shimadzu; HP-INOWAX capillary column, 30 m, 0.25 mm, Agilent).

### Statistics

Statistics used for each experiment was described in Figure Legends. Briefly, *p* values were calculated by using two-tailed Student t-test for embryonic lethality, body size, RNA-seq., qRT-PCR, fluorescence imaging, Oil red O, and GC/MS assays. For survival assays, statistical analysis was performed by using OASIS 2 (online application of survival analysis, http://sbi.postech.ac.kr/oasis2) (Han et al., 2016) and *p* values were calculated by using log-rank (Mantel-Cox method) test. Significance of the overlaps among gene sets was calculated by using hypergeometric probability test (http://systems.crump.ucla.edu/hypergeometric/index.php, provided by Graeber lab).

## Acknowledgments

We thank all Lee laboratory members for help and discussion. We also thank Craig Mello lab for providing CRISPR/Cas9 protocol, and Marc Pilon and Junho Lee labs for strains. This work was supported by the Korean Government (MSIP) through the National Research Foundation of Korea (NRF) (NRF-2016R1E1A1A01941152) to S-J. V. L. and by Canadian Institutes of Health Research (CIHR) (PJT-153199) to S.T.

## Author contributions

D.L. contributed to designing and performing the majority of experiments for all the assays described in the manuscript, data analysis, and writing manuscript; S.W.A.A. and Y.J. contributed to survival assay experiments; Y.Y. and Y.L. contributed to lipid composition analysis; Y.R. and C.M.H. contributed to confocal microscopy imaging; D.M. contributed to RNA-seq. analysis; G.Y.S.G, A.B., and S.T. contributed to reproduction assay experiments; S.-J.V.L. contributed to designing all experiments, data analysis, and writing manuscript.

## References

Allen, B.L., and Taatjes, D.J. 2015. The Mediator complex: a central integrator of transcription. Nature reviews Molecular cell biology. doi: 10.1038/nrm3951

Artan, M., Jeong, D.E., Lee, D., Kim, Y.I., Son, H.G., Husain, Z., Kim, J., Altintas, O., Kim, K., Alcedo, J., et al. 2016. Food-derived sensory cues modulate longevity via distinct neuroendocrine insulin-like peptides. Genes & development. doi: 10.1101/gad.279448.116

Brock, T.J., Browse, J., and Watts, J.L. 2007. Fatty acid desaturation and the regulation of adiposity in *Caenorhabditis elegans*. Genetics. doi: 10.1534/genetics.107.071860

Brokate-Llanos, A.M., Garzon, A., and Munoz, M.J. 2014. *Escherichia coli* carbon source metabolism affects longevity of its predator *Caenorhabditis elegans*. Mechanisms of ageing and development. doi: 10.1016/j.mad.2014.09.001

Brunquell, J., Morris, S., Lu, Y., Cheng, F., and Westerheide, S.D. 2016. The genome-wide role of HSF-1 in the regulation of gene expression in *Caenorhabditis elegans*. BMC genomics. doi: 10.1186/s12864-016-2837-5

Chen, Y.C., Chen, H.J., Tseng, W.C., Hsu, J.M., Huang, T.T., Chen, C.H., and Pan, C.L. 2016. A *C. elegans* Thermosensory Circuit Regulates Longevity through *crh*-*1*/CREB-Dependent *flp*-*6* Neuropeptide Signaling. Developmental cell. doi: 10.1016/j.devcel.2016.08.021

Chi, Y., and Gupta, R.K. 1998. Alterations in membrane fatty acid unsaturation and chain length in hypertension as observed by 1H NMR spectroscopy. American journal of hypertension. doi: Conti, B. 2008. Considerations on temperature, longevity and aging. Cellular and molecular life sciences : CMLS. doi: 10.1007/s00018-008-7536-1

Conti, B., Sanchez-Alavez, M., Winsky-Sommerer, R., Morale, M.C., Lucero, J., Brownell, S., Fabre, V., Huitron-Resendiz, S., Henriksen, S., Zorrilla, E.P., et al. 2006. Transgenic mice with a reduced core body temperature have an increased life span. Science (New York, NY). doi: 10.1126/science.1132191

de Hoon, M.J., Imoto, S., Nolan, J., and Miyano, S. 2004. Open source clustering software. Bioinformatics (Oxford, England). doi: 10.1093/bioinformatics/bth078

Deline, M.L., Vrablik, T.L., and Watts, J.L. 2013. Dietary supplementation of polyunsaturated fatty acids in *Caenorhabditis elegans*. Journal of visualized experiments : JoVE. doi: 10.3791/50879

Devkota, R., Svensk, E., Ruiz, M., Stahlman, M., Boren, J., and Pilon, M. 2017. The adiponectin receptor AdipoR2 and its *Caenorhabditis elegans* homolog PAQR-2 prevent membrane rigidification by exogenous saturated fatty acids. PLoS genetics. doi: 10.1371/journal.pgen.1007004

Goh, G.Y., Martelli, K.L., Parhar, K.S., Kwong, A.W., Wong, M.A., Mah, A., Hou, N.S., and Taubert, S. 2014. The conserved Mediator subunit MDT-15 is required for oxidative stress responses in *Caenorhabditis elegans*. Aging cell. doi: 10.1111/acel.12154

Grants, J.M., Goh, G.Y., and Taubert, S. 2015. The Mediator complex of *Caenorhabditis elegans:* insights into the developmental and physiological roles of a conserved transcriptional coregulator. Nucleic acids research. doi: 10.1093/nar/gkv037

Han, S., Schroeder, E.A., Silva-Garcia, C.G., Hebestreit, K., Mair, W.B., and Brunet, A. 2017. Mono-unsaturated fatty acids link H3K4me3 modifiers to *C. elegans* lifespan. Nature. doi: 10.1038/nature21686

Han, S.K., Lee, D., Lee, H., Kim, D., Son, H.G., Yang, J.S., Lee, S.V., and Kim, S. 2016. OASIS 2: online application for survival analysis 2 with features for the analysis of maximal lifespan and healthspan in aging research. Oncotarget. doi: 10.18632/oncotarget.11269

Heintz, C., Doktor, T.K., Lanjuin, A., Escoubas, C., Zhang, Y., Weir, H.J., Dutta, S., Silva-Garcia, C.G., Bruun, G.H., Morantte, I., et al. 2017. Splicing factor 1 modulates dietary restriction and TORC1 pathway longevity in *C. elegans*. Nature. doi: 10.1038/nature20789

Higuchi-Sanabria, R., Frankino, P.A., Paul, J.W., 3rd, Tronnes, S.U., and Dillin, A. 2018. A Futile Battle? Protein Quality Control and the Stress of Aging. Developmental cell. doi: 10.1016/j.devcel.2017.12.020

Holthuis, J.C., and Menon, A.K. 2014. Lipid landscapes and pipelines in membrane homeostasis. Nature. doi: 10.1038/nature13474

Horikawa, M., Sural, S., Hsu, A.L., and Antebi, A. 2015. Co-chaperone p23 regulates *C. elegans* Lifespan in Response to Temperature. PLoS genetics. doi: 10.1371/journal.pgen.1005023

Huang da, W., Sherman, B.T., and Lempicki, R.A. 2009. Systematic and integrative analysis of large gene lists using DAVID bioinformatics resources. Nature protocols. doi: 10.1038/nprot.2008.211

Jeong, D.E., Artan, M., Seo, K., and Lee, S.J. 2012. Regulation of lifespan by chemosensory and thermosensory systems: findings in invertebrates and their implications in mammalian aging. Frontiers in genetics. doi: 10.3389/fgene.2012.00218

Kim, H.E., Grant, A.R., Simic, M.S., Kohnz, R.A., Nomura, D.K., Durieux, J., Riera, C.E., Sanchez, M., Kapernick, E., Wolff, S., et al. 2016. Lipid Biosynthesis Coordinates a Mitochondrial-to-Cytosolic Stress Response. Cell. doi: 10.1016/j.cell.2016.08.027

Labbadia, J., Brielmann, R.M., Neto, M.F., Lin, Y.F., Haynes, C.M., and Morimoto, R.I. 2017. Mitochondrial Stress Restores the Heat Shock Response and Prevents Proteostasis Collapse during Aging. Cell reports. doi: 10.1016/j.celrep.2017.10.038

Lee, D., Jeong, D.E., Son, H.G., Yamaoka, Y., Kim, H., Seo, K., Khan, A.A., Roh, T.Y., Moon, D.W., Lee, Y., et al. 2015. SREBP and MDT-15 protect *C. elegans* from glucose-induced accelerated aging by preventing accumulation of saturated fat. Genes & development. doi: 10.1101/gad.266304.115

Lee, S.J., Hwang, A.B., and Kenyon, C. 2010. Inhibition of respiration extends *C. elegans* life span via reactive oxygen species that increase HIF-1 activity. Current biology : CB. doi: 10.1016/j.cub.2010.10.057

Lee, S.J., and Kenyon, C. 2009. Regulation of the longevity response to temperature by thermosensory neurons in *Caenorhabditis elegans*. Current biology : CB. doi: 10.1016/j.cub.2009.03.041

Lee, S.J., Murphy, C.T., and Kenyon, C. 2009. Glucose shortens the life span of *C. elegans* by downregulating DAF-16/FOXO activity and aquaporin gene expression. Cell metabolism. doi: 10.1016/j.cmet.2009.10.003

Lopez-Otin, C., Blasco, M.A., Partridge, L., Serrano, M., and Kroemer, G. 2013. The hallmarks of aging. Cell. doi: 10.1016/j cell.2013.05.039

Ma, D.K., Li, Z., Lu, A.Y., Sun, F., Chen, S., Rothe, M., Menzel, R., Sun, F., and Horvitz, H.R. 2015. Acyl-CoA Dehydrogenase Drives Heat Adaptation by Sequestering Fatty Acids. Cell. doi: 10.1016/j.cell.2015.04.026

McKay, R.M., McKay, J.P., Avery, L., and Graff, J.M. 2003. *C elegans:* a model for exploring the genetics of fat storage. Developmental cell. doi: 10.1016/S1534-5807(02)00411-2

Morley, J.F., Brignull, H.R., Weyers, J.J., and Morimoto, R.I. 2002. The threshold for polyglutamine-expansion protein aggregation and cellular toxicity is dynamic and influenced by aging in *Caenorhabditis elegans*. Proceedings of the National Academy of Sciences of the United States of America. doi: 10.1073/pnas.152161099

Murphy, C.T., McCarroll, S.A., Bargmann, C.I., Fraser, A., Kamath, R.S., Ahringer, J., Li, H., and Kenyon, C. 2003. Genes that act downstream of DAF-16 to influence the lifespan of *Caenorhabditis elegans*. Nature. doi: 10.1038/nature01789

Murray, P., Hayward, S.A., Govan, G.G., Gracey, A.Y., and Cossins, A.R. 2007. An explicit test of the phospholipid saturation hypothesis of acquired cold tolerance in *Caenorhabditis elegans*. Proceedings of the National Academy of Sciences of the United States of America. doi: 10.1073/pnas.0609590104

O’Rourke, E.J., Kuballa, P., Xavier, R., and Ruvkun, G. 2013. omega-6 Polyunsaturated fatty acids extend life span through the activation of autophagy. Genes & development. doi: 10.1101/gad.205294.112

O’Rourke, E.J., Soukas, A.A., Carr, C.E., and Ruvkun, G. 2009. *C. elegans* major fats are stored in vesicles distinct from lysosome-related organelles. Cell metabolism. doi: 10.1016/j.cmet.2009.10.002

Pang, S., Lynn, D.A., Lo, J.Y., Paek, J., and Curran, S.P. 2014. SKN-1 and Nrf2 couples proline catabolism with lipid metabolism during nutrient deprivation. Nature communications. doi: 10.1038/ncomms6048

Pertea, M., Kim, D., Pertea, G.M., Leek, J.T., and Salzberg, S.L. 2016. Transcript-level expression analysis of RNA-seq experiments with HISAT, StringTie and Ballgown. Nature protocols. doi: 10.1038/nprot.2016.095

Pukkila-Worley, R., Feinbaum, R.L., McEwan, D.L., Conery, A.L., and Ausubel, F.M. 2014. The evolutionarily conserved mediator subunit MDT-15/MED15 links protective innate immune responses and xenobiotic detoxification. PLoS pathogens. doi: 10.1371/journal.ppat.1004143

Rea, S.L., Ventura, N., and Johnson, T.E. 2007. Relationship between mitochondrial electron transport chain dysfunction, development, and life extension in *Caenorhabditis elegans*. PLoS biology. doi: 10.1371/journal.pbio.0050259

Ritchie, M.E., Phipson, B., Wu, D., Hu, Y., Law, C.W., Shi, W., and Smyth, G.K. 2015. limma powers differential expression analyses for RNA-sequencing and microarray studies. Nucleic acids research. doi: 10.1093/nar/gkv007

Robinson, M.D., McCarthy, D.J., and Smyth, G.K. 2010. edgeR: a Bioconductor package for differential expression analysis of digital gene expression data. Bioinformatics (Oxford, England). doi: 10.1093/bioinformatics/btp616

Saldanha, A.J. 2004. Java Treeview‐‐extensible visualization of microarray data. Bioinformatics (Oxford, England). doi: 10.1093/bioinformatics/bth349

Schleit, J., Wall, V.Z., Simko, M., and Kaeberlein, M. 2011. The MDT-15 subunit of mediator interacts with dietary restriction to modulate longevity and fluoranthene toxicity in *Caenorhabditis elegans*. PloS one. doi: 10.1371/journal.pone.0028036

Schneider, C.A., Rasband, W.S., and Eliceiri, K.W. 2012. NIH Image to ImageJ: 25 years of image analysis. Nature methods. doi: 10.1038/nmeth.2089

Seo, K., Choi, E., Lee, D., Jeong, D.E., Jang, S.K., and Lee, S.J. 2013. Heat shock factor 1 mediates the longevity conferred by inhibition of TOR and insulin/IGF-1 signaling pathways in *C. elegans*. Aging cell. doi: 10.1111/acel.12140

Soderberg, M., Edlund, C., Kristensson, K., and Dallner, G. 1991. Fatty acid composition of brain phospholipids in aging and in Alzheimer’s disease. Lipids. doi:

Son, H.G., and Lee, S.V. 2017. Longevity regulation by NMD-mediated mRNA quality control. BMB reports. doi: 10.5483/BMBRep.2017.50.4.045

Son, H.G., Seo, M., Ham, S., Hwang, W., Lee, D., An, S.W., Artan, M., Seo, K., Kaletsky, R., Arey, R.N., et al. 2017. RNA surveillance via nonsense-mediated mRNA decay is crucial for longevity in *daf*-*2*/insulin/IGF-1 mutant *C. elegans*. Nature communications. doi: 10.1038/ncomms14749

Steinbaugh, M.J., Narasimhan, S.D., Robida-Stubbs, S., Moronetti Mazzeo, L.E., Dreyfuss, J.M., Hourihan, J.M., Raghavan, P., Operana, T.N., Esmaillie, R., and Blackwell, T.K. 2015. Lipid-mediated regulation of SKN-1/Nrf in response to germ cell absence. eLife. doi: 10.7554/eLife.07836

Stiernagle, T. 2006. Maintenance of *C. elegans*. WormBook : the online review of C elegans biology. doi: 10.1895/wormbook.1.101.1

Svensk, E., Devkota, R., Stahlman, M., Ranji, P., Rauthan, M., Magnusson, F., Hammarsten, S., Johansson, M., Boren, J., and Pilon, M. 2016. *Caenorhabditis elegans* PAQR-2 and IGLR-2 Protect against Glucose Toxicity by Modulating Membrane Lipid Composition. PLoS genetics. doi: 10.1371/journal.pgen.1005982

Svensk, E., Stahlman, M., Andersson, C.H., Johansson, M., Boren, J., and Pilon, M. 2013. PAQR-2 regulates fatty acid desaturation during cold adaptation in *C. elegans*. PLoS genetics. doi: 10.1371/journal.pgen.1003801

Sztolsztener, M.E., Dobrzyn, A., Pikula, S., Tylki-Szymanska, A., and Bandorowicz-Pikula, J. 2012. Impaired dynamics of the late endosome/lysosome compartment in human Niemann-Pick type C skin fibroblasts carrying mutation in NPC1 gene. Molecular bioSystems. doi: 10.1039/c2mb05447g

Tabrez, S.S., Sharma, R.D., Jain, V., Siddiqui, A.A., and Mukhopadhyay, A. 2017. Differential alternative splicing coupled to nonsense-mediated decay of mRNA ensures dietary restriction-induced longevity. Nature communications. doi: 10.1038/s41467-017-00370-5

Tanaka, T., Ikita, K., Ashida, T., Motoyama, Y., Yamaguchi, Y., and Satouchi, K. 1996. Effects of growth temperature on the fatty acid composition of the free-living nematode *Caenorhabditis elegans*. Lipids. doi: 10.1007/BF02524292

Taubert, S., Hansen, M., Van Gilst, M.R., Cooper, S.B., and Yamamoto, K.R. 2008. The Mediator subunit MDT-15 confers metabolic adaptation to ingested material. PLoS genetics. doi: 10.1371/journal.pgen.1000021

Taubert, S., Van Gilst, M.R., Hansen, M., and Yamamoto, K.R. 2006. A Mediator subunit, MDT-15, integrates regulation of fatty acid metabolism by NHR-49-dependent and -independent pathways in *C. elegans*. Genes & development. doi: 10.1101/gad.1395406

Van Gilst, M.R., Hadjivassiliou, H., Jolly, A., and Yamamoto, K.R. 2005. Nuclear hormone receptor NHR-49 controls fat consumption and fatty acid composition in *C. elegans*. PLoS biology. doi: 10.1371/journal.pbio.0030053

Wang, L., Folsom, A.R., and Eckfeldt, J.H. 2003. Plasma fatty acid composition and incidence of coronary heart disease in middle aged adults: the Atherosclerosis Risk in Communities (ARIC) Study. Nutrition, metabolism, and cardiovascular diseases : NMCD. doi: 10.1016/S0939-4753(03)80029-7

Xiao, R., Zhang, B., Dong, Y., Gong, J., Xu, T., Liu, J., and Xu, X.Z. 2013. A genetic program promotes *C. elegans* longevity at cold temperatures via a thermosensitive TRP channel. Cell. doi: 10.1016/j.cell.2013.01.020

Yang, F., Vought, B.W., Satterlee, J.S., Walker, A.K., Jim Sun, Z.Y., Watts, J.L., DeBeaumont, R., Saito, R.M., Hyberts, S.G., Yang, S., et al. 2006. An ARC/Mediator subunit required for SREBP control of cholesterol and lipid homeostasis. Nature. doi: 10.1038/nature04942

Yu, Y, Mutlu, A.S., Liu, H., and Wang, M.C. 2017. High-throughput screens using photo-highlighting discover BMP signaling in mitochondrial lipid oxidation. Nature communications. doi: 10.1038/s41467-017-00944-3

Zhang, L., Ward, J.D., Cheng, Z., and Dernburg, A.F. 2015. The auxin-inducible degradation (AID) system enables versatile conditional protein depletion in *C. elegans*. Development (Cambridge, England). doi: 10.1242/dev.129635

